# Regulation of a nickel tolerance operon conserved in *Mesorhizobium* strains from serpentine soils

**DOI:** 10.1101/2025.07.11.664438

**Authors:** Kyson T. Jensen, Joel S. Griffitts

## Abstract

The *Mesorhizobium nre* (nickel resistance) operon has previously been shown to mediate Ni tolerance in serpentine soils with naturally high concentrations of this transition metal. In most serpentine-derived strains evaluated, the putative efflux genes *nreX* and *nreY* are conserved, along with a small gene (*nreA*) encoding a CsoR/RcnR-family transcriptional regulator. CsoR/RcnR-family regulators are small (around 100 amino acids in length), they bind transition metals, and they use an unconventional and poorly understood DNA-binding mechanism. NreA is 93 amino acids in length and belongs to a poorly characterized clade within the CsoR/RcnR family. This investigation is focused on regulatory DNA elements that functionally interact with *Mesorhizobium* NreA, as well as amino acid residues in NreA that influence its regulatory activity. We show that NreA is a transcriptional repressor that is responsive to exogenous Ni. The Ni-responsive promoter is immediately upstream of the *nreAXY* operon, resulting in a leaderless transcript. A partially palindromic operator sequence occupies the spacer region between the −35 and −10 promoter elements. Mutational analysis of the operator highlights functionally crucial base pairs occupying opposite ends of the palindromic operator. Changes to conserved residues in the NreA polypeptide result in varying effects, with changes predicted to prevent Ni binding leading to a super-repressor phenotype. Structural modeling of the NreA-operator complex provides a plausible mechanism for DNA binding by a tetrameric form of NreA, with DNA contact sites along a positively charged surface, and the modeled contact sites agree well with the mutational analysis.

**IMPORTANCE:** Bacteria employ diverse mechanisms for maintaining optimal intracellular levels of bioactive metals. Isolates of *Mesorhizobium* bacteria from nickel-rich serpentine soils possess a small genetic region controlling Ni efflux. This region is subject to transcriptional regulation via the small repressor protein NreA. In this work, the essential components of the NreA protein and the DNA operator with which it interacts are defined, enabling potential adaptation of this system for metal sensing or other technologies requiring a compact inducible gene expression module.

## INTRODUCTION

Bacteria have evolved diverse mechanisms to cope with metal stress, including efflux, sequestration, and detoxification of toxic metal ions (1–3). The fitness costs associated with these systems, and the need to precisely regulate the balance of intracellular metals, requires bacteria to maintain precise control of metal stress responses (4). This control is primarily implemented through metal-responsive transcription factors, or metalloregulators (5). Metalloregulators, which may be transcriptional activators or repressors, are generally homo-oligomeric proteins that interact with DNA operators in upstream regulatory regions for genes controlling metal homeostasis. Metalloregulators also interact with specific metal ions that modulate binding to cognate operators (1, 2).

Repressors from the CsoR/RcnR metalloregulator family have received attention due to their small size and poorly defined mechanism of DNA binding (6–8). CsoR is a copper-responsive regulator used by various Gram-positive bacteria, and RcnR is a nickel- and cobalt-responsive regulator found in Gram-negative enteric bacteria including *E. coli* (9, 10). Crystal structures of CsoR from *Mycobacterium tuberculosis* and *Streptococcus lividans* reveal a homodimeric or homotetrameric complex of alpha-helical bundles in which metal ions are coordinated in a cysteine- and histidine-containing coordination site formed in an intersubunit bridge (9, 11). A DNA-bound structure for this family has not been reported, though it has been presumed that a face of the tetrameric structure with strong positive electrostatic potential may mediate DNA binding (6, 8, 12, 13). Copper binding drives CsoR disengagement from operator DNA, and alterations to CsoR that prevent metal binding cause constitutive repression of the regulated operon (9, 14, 15). Metal coordination sites in CsoR and RcnR family members appear to be located in analogous intersubunit positions, though copper binding to CsoR is mediated by a Cys2His1 motif, while nickel/cobalt binding to RcnR is mediated by a Cys1His3 motif (9, 16).

A recent study of *Mesorhizobium* isolates from nickel-rich serpentine soils pointed us to a conserved nickel resistance operon containing a gene for a CsoR/RcnR-like protein that we termed NreA (17). The *nreA* gene is the most upstream in an operon that also encodes two independently functioning nickel efflux transporters: NreX and NreY (17). Here we characterize the mechanism by which NreA transcriptionally regulates the *nreAXY* operon.

## RESULTS

### Conserved sequences identify the minimal *nreAXY* promoter/operator

*Mesorhizobium* strain C089B is derived from a serpentine soil plot at McLaughlin natural reserve and contains a *nreAXY* operon (Fig. 1A). The NreA polypeptide is similar in sequence to members of the CsoR/RcnR metalloregulator family, though it fits best into a phylogenetic clade with other uncharacterized NreA-like proteins from alpha- and beta-proteobacteria (Fig. 1B). In an initial attempt to discern a *cis*-acting regulatory region upstream of *nreA* that may be controlled by NreA, we compared sequences from 21 additional serpentine soil-derived *Mesorhizobium* strains (Fig. 1C). This DNA sequence alignment encompassed the *nreA* start codon on the right, and approximately 65 bp upstream of the start codon. The sequence logo generated in Fig. 1C highlights a strongly conserved TTG motif most likely constituting the −35 RNA polymerase (RNAP) binding motif. A TAGNAT motif 20 bp to the right of this is likely a corresponding −10 RNAP binding motif. There is a strongly conserved, partially palindromic sequence positioned between these putative −35 and −10 elements (the spacer region), which we presumed may function as an NreA operator (see Fig. 1C). Notably, these elements occur quite close to the *nreA* start codon, suggesting they control transcription of an mRNA devoid of any substantial 5’ untranslated RNA.

**FIG 1.**
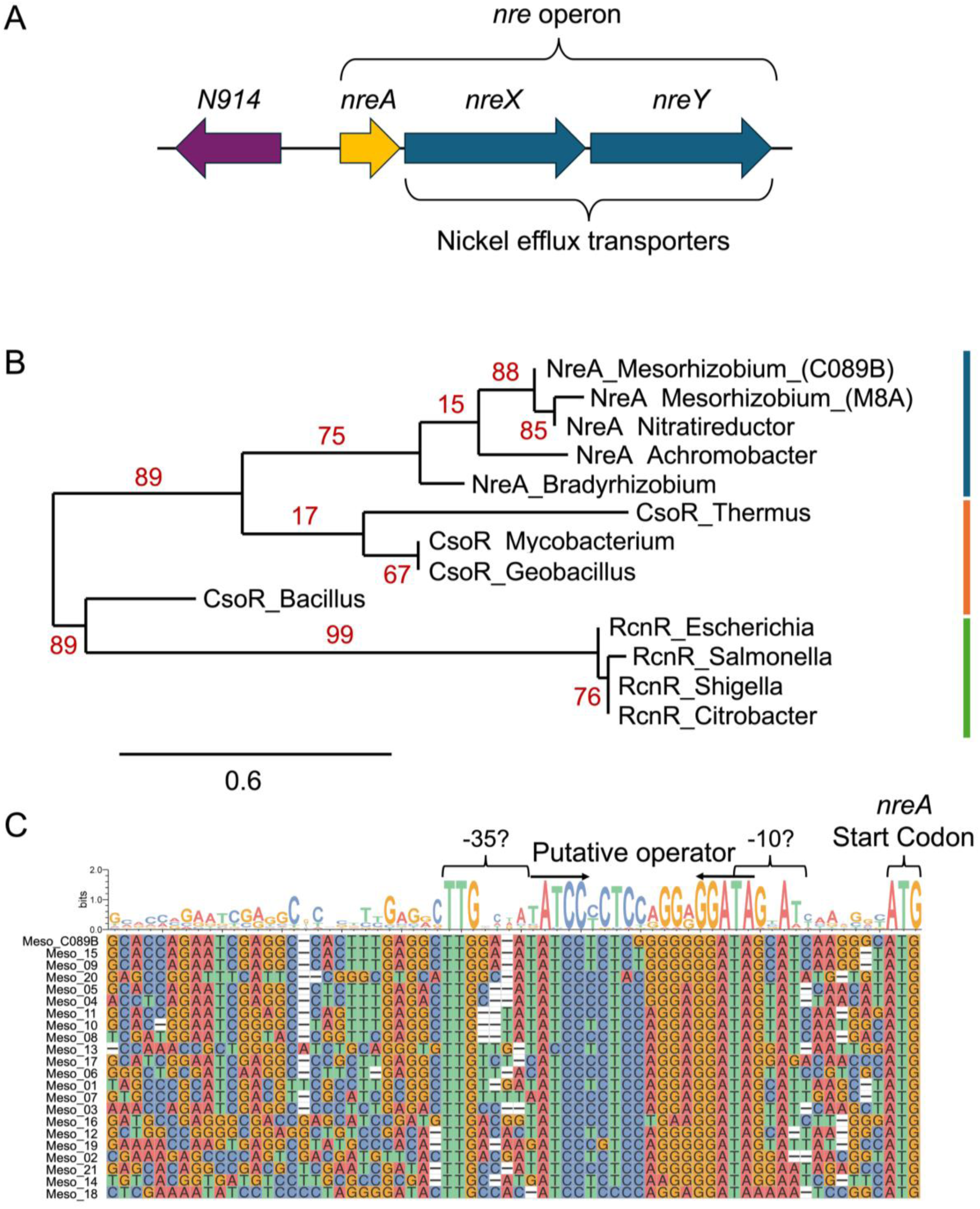
Comparison of *nreA* upstream regions points to probable regulatory elements. (A) Map of the *nre* operon identified in *Mesorhizobium* strain C089B. (B) Protein phylogenetic tree of members of the CsoR/RcnR family of transcriptional regulators. Support values are in red and the scale bar indicates substitutions per site; NreA, CsoR, and RcnR clades are represented with blue, orange, and green lines, respectively. (C) Multiple sequence alignment of the *nre* upstream region from 21 different *Mesorhizobium* isolates, including C089B. Proposed promoter/operator elements are labeled, with arrows showing palindromic regions.

The conserved upstream sequence elements in the alignment in Fig. 1C set the stage for plasmid-based *lacZ* reporter analyses. We probed the functionality of a 120-bp upstream region, or truncated variants thereof (Fig. 2A). Reporter assays were facilitated by a broad-host range shuttle vector (pKJ138; Fig. 2B) containing *lacZ* with a synthetic ribosome-binding site. This vector can accommodate the presence or absence of *nreA* downstream of the constitutively expressed kanamycin resistance gene. In this way, *nreA* expression is uncoupled from the transcriptional control region under investigation (see Fig. 2B). Most of the reporter gene tests described here were carried out in *Agrobacterium fabrum*, owing to its faster growth and easier genetic manipulation compared to wild *Mesorhizobium* strains. In this heterologous species, we could also be more confident in the sufficiency of the genetic elements contained in these plasmids to carry out the observed regulatory outcomes.

**FIG 2.**
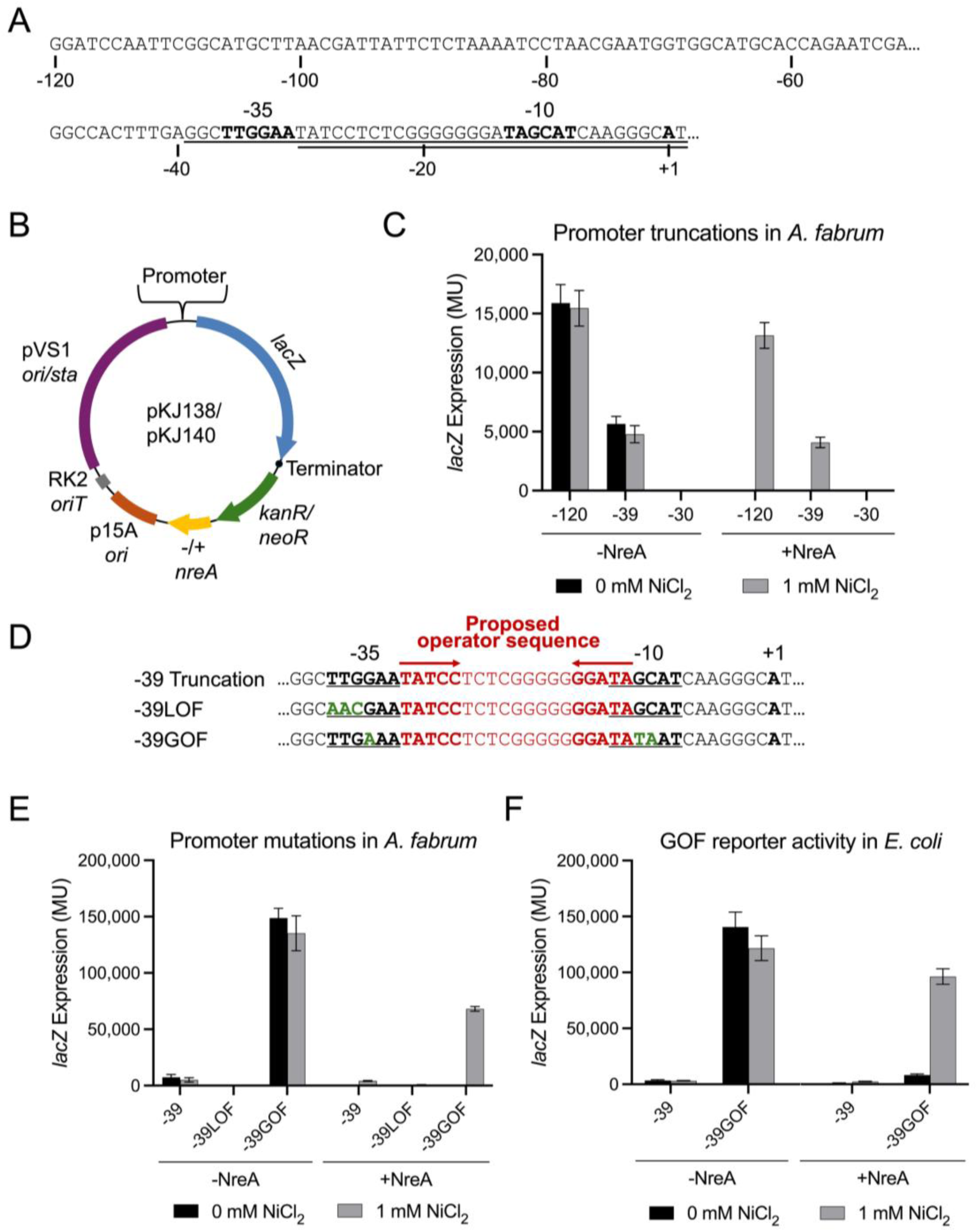
Identification of a minimal *nre* promoter and its functional elements. (A) Upstream sequence of the *nre* operon. Underlined sequences show −39 and −30 truncations. Bolded characters represent proposed −35 element, −10 element, and the translational start site. (B) Map of the binary plasmid used to test promoter truncations and mutations in either *A. fabrum* or *E. coli*. (C) Reporter gene activity from truncations of the *nre* promoter region. *A. fabrum* cells harboring *lacZ* fusions with or without *nreA* were grown in the presence or absence of supplemental NiCl_2_, and β-galactosidase activity was measured in Miller units (MU). Error bars represent standard deviation from the mean (n=6). (D) Sequence alignment showing the −39LOF and −39GOF mutations (green). Underlined are putative −35 and −10 elements. Red denotes the putative operator sequence and inverted arrows indicate the palindromic regions within this sequence. (E,F) Functional comparisons of promoter mutants shown in (D) carried out in both *A. fabrum* (E) and *E. coli* (F). Data are formatted as in (C). Error bars represent standard deviation elements, we intentionally avoided changes to the putative repressor-binding operator comprising the −30 to −12 region (red in Fig. 2D).

When the full 120-bp region is tested in this system, we observe high-level expression that is repressed by *nreA* co-expression, and this *nreA*-dependent repression is alleviated by 1 mM NiCl_2_ (Fig. 2C). This NiCl_2_ concentration was chosen because it is the highest concentration that allows robust growth of *A. fabrum*. We next tested whether truncation of the promoter region to the −39 position would still support Ni-inducible expression, supposing that maintenance of the core −35 and −10 promoter elements, as well as the putative operator site, would allow regulated expression similar to the full-length 120-bp segment. The −39 construct supports reporter gene expression that is repressed by NreA and induced by NiCl_2_, but induced expression levels are lower by approximately 3-fold compared to the full-length construct (Fig. 2C). This may be explained by a positive regulatory element existing somewhere in the −120 to −39 region; however, given the robust inducible response we observe from the −39 construct, we proceeded to characterize the regulatory features within this much shorter sequence. As expected, the −30 construct did not support reporter gene expression under any condition, presumably due to loss of the −35 TTG element (Fig. 2C). In further support of the role of the −35 TTG element as a positive regulatory element, mutation of this sequence to AAC within the context of the −39 construct (−39LOF, Fig. 2D) yielded undetectable expression (Fig. 2E). On the other hand, a 3-bp change to the −39 construct designed to bring both the −35 and −10 elements closer to a prokaryotic promoter consensus sequence (−39GOF, Fig. 2D) led to greatly increased expression while maintaining the NreA-dependent, Ni-inducible property (Fig. 2E). Under inducing conditions, −39GOF supports approximately 16-fold higher expression, more than compensating for loss of the transcriptionally enhancing activity present in the −120 to −39 region. The - 39GOF-*lacZ* construct also supports expression in the more distantly related *E. coli*, and this expression is also NreA-repressed and Ni-induced (Fig. 2F). It should be noted that in our attempt to enhance these positive regulatory

### The *nreAXY* operon is expressed as a leaderless mRNA

The close proximity of the proposed promoter elements to the *nreA* start codon suggests that the transcript may be leaderless or may contain only a few untranslated nucleotides. Sequencing results from 5’ rapid amplification of cDNA ends (5’-RACE) supports production of a leaderless transcript (Fig. 3). This test was carried out using an *A. fabrum* strain carrying the P*nreA*-*lacZ* - 120 construct described above. This reporter fusion includes the first two base pairs of the native *nreA* start codon fused to a synthetic ribosome binding site for efficient translation of the *lacZ* coding sequence, as illustrated in Fig. 3. In this context, the mRNA 5’ end mapped to the A nucleotide that corresponds to the native *nreA* start codon. Unfortunately, difficulty in isolating quality RNA from *Mesorhizobium* cells prevented this test from being carried out in the strain naturally harboring the *nreAXY* operon. This finding defines the +1 site of transcription, as well as confirming the locations of the −10 and −35 promoter elements proposed above.

**FIG 3.**
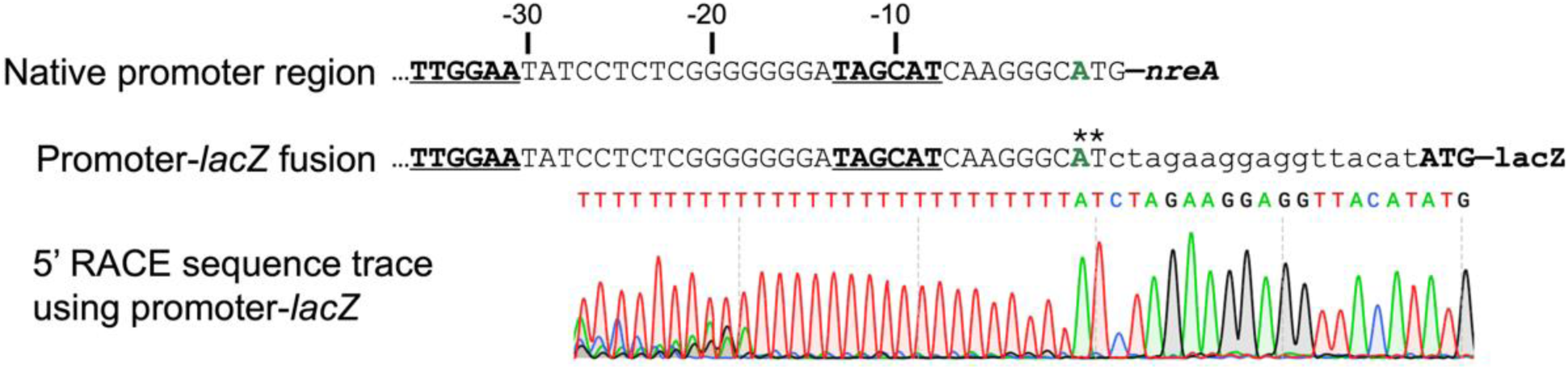
Identification of the *nre* transcription start site by 5’-RACE. Analysis was carried out in *A. fabrum* carrying a promoter-*lacZ* fusion that includes the first two nucleotides of the native *nreA* start codon (**). Underlined are the −35 and −10 elements of the *nre* promoter. Green is the first nucleotide of the native *nreA* start codon. Bottom, inverted bottom-strand sequence trace acquired after poly-A tailing of the reverse-transcription product and subsequent PCR amplification.

### Palindromic ends of the *nre* operator sequence are vital for repression

We next evaluated the role of the −30 to −12 spacer sequence, which we presumed to be the site of negative regulation by NreA. A set of mutant reporter fusions constituting a “scan” of AAA substitutions was tested (Fig. 4), and expression values indicate that the distal palindromic ends of this spacer (i.e. TCCN_9_GGA) are particularly crucial for negative regulation: both the TCC->AAA (*o*_mut1) and the GGA->AAA (*o*_mut5) substitutions result in fully constitutive reporter gene expression. The three AAA substitutions across the central N_9_ region (*o*_mut2 through *o*_mut4) resulted in only weakly constitutive phenotypes compared to the wild-type operator (*o*_wt_). In other words, these AAA mutants are mostly inducible (see Fig. 4B).

**FIG 4.**
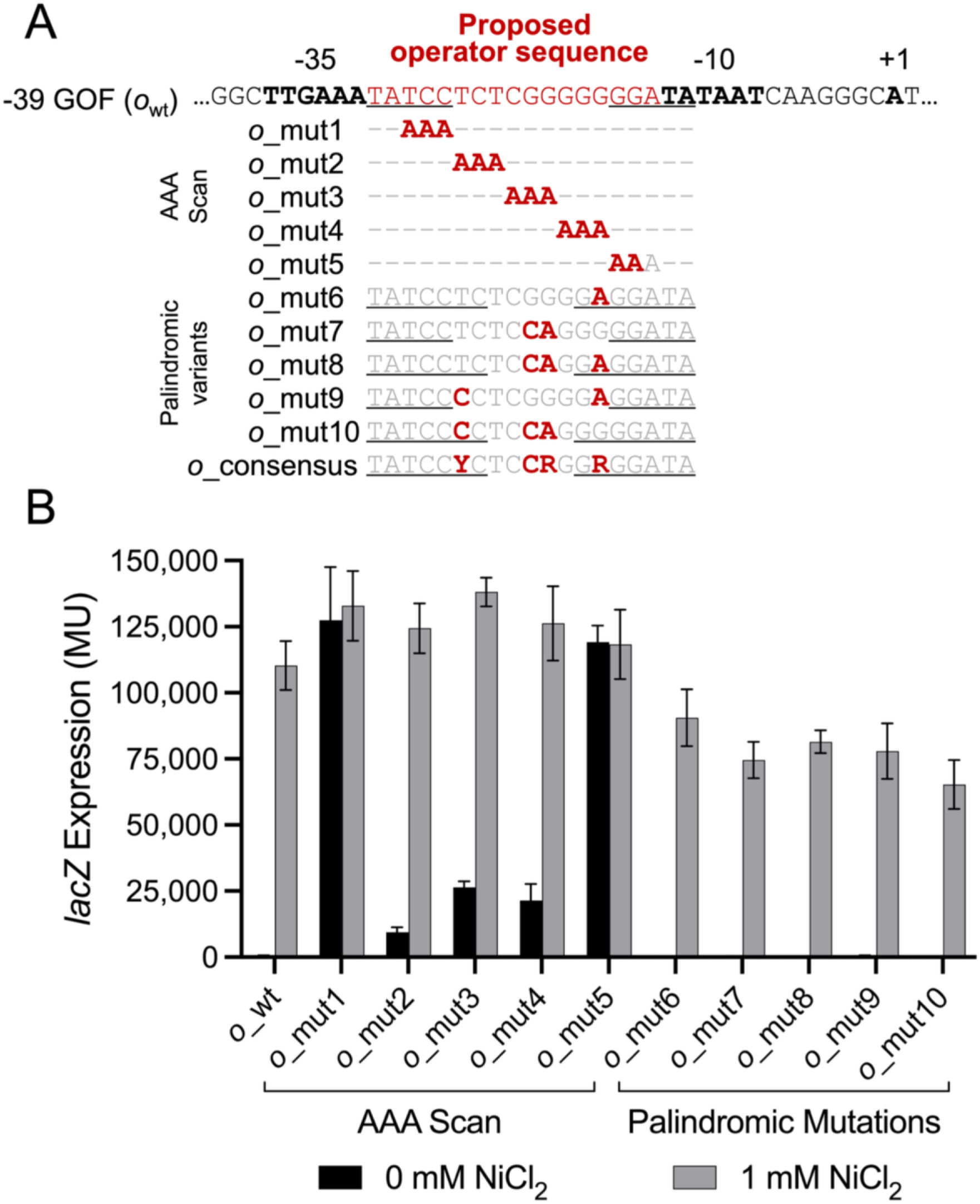
Identification of operator elements essential for NreA-mediated regulation. (A) Mutational analysis of the operator site. Black/bolded: −35 and −10 elements. Red/bold: mutations. Black underlines: palindromic regions. (B) Reporter gene activity resulting from mutations of the *nre* operator sequence. *A. fabrum* cells (*nreA^+^*) harboring *lacZ* fusions were grown in the presence or absence of supplemental NiCl_2_, and β-galactosidase activity was measured in Miller units (MU). Error bars represent the standard deviation from the mean (n=6).

The TCCN_9_GGA operator sequence from *Mesorhizobium* strain C089B (*o*_wt_) is partially palindromic, and we know from results just presented that the distal ends of this palindrome are particularly crucial for operator function. The consensus sequence derived from an alignment of 22 *Mesorhizobium nre* promoter/operator regions (Fig. 1C) is more extensively palindromic, with the sequence TATCCYCtccrgGRGGATA, and some strains such as C089B contain a particularly G-rich sequence through the central region. It has been proposed for the metalloregulators RcnR and CsoR that a similar G-rich operator sequence may bring the DNA into an alternative hybrid A/B-form to facilitate repressor binding (15, 18). In Fig. 4, operator variants *o_*mut6 through *o*_mut10 represent various alternative sequences that either increase the palindromic symmetry of the central region or lessen its G-richness. In all cases, the variant operators support robust NreA-dependent repression and Ni inducibility. That one of these variants (*o*_mut8) has considerably less G-rich content and likely does not take on the alternative A/B hybrid DNA configuration suggests that NreA is able to function on operators composed of ordinary B-form DNA. Given that operator *o*_mut6 through *o*_mut10 function in reporter gene repression, we investigated whether any of these support even stronger repression than the native operator, *o*_wt_. We explored this low-range (leaky) expression by adding cells cultured overnight in the absence of Ni to our standard Miller assay and incubating the reaction for an extended period. Operators *o*_mut7 and *o*_mut10 showed 3 times more repression than *o*_wt_, with *o*_mut8 being intermediate in repression (Fig. S1). These three clones share the GG->CA change in the middle of the operator, suggesting that this specific change leads to stronger repressor binding. Operator variants *o*_mut6 and *o*_mut9, which do not contain the GG->CA substitutions, yield levels of repression somewhat worse than *o*_wt_.

### Conserved NreA residues influence either DNA-binding or Metal-binding activities

The *Mesorhizobium* NreA protein bears extensive sequence similarity to CsoR/RcnR family members (see sequence alignment in Fig. 5A). In the structural model of NreA as a tetramer (Fig. 5B-C), the amino acids best conserved in the sequence alignment generally correspond either to an intersubunit metal-binding pocket (NreA residues C38, H63, H66, and C67), or to a potential DNA-binding surface (NreA residues R17, R20, Q44, and K91). We performed inducibility tests for strains in which each of these conserved residues was substituted with alanine. In Fig. 5D, we see that alterations across the putative DNA binding surface yield mostly or completely constitutive phenotypes, consistent with loss of operator binding. Substitutions around the putative Ni-binding pocket, on the other hand, give rise to mostly or completely super-repressing phenotypes–that is, repression persists even in the presence of Ni. The partial functionality of the H66A variant is consistent with the structural model (Fig. 5C), which places this residue at a location peripheral to the binding pocket. On the other hand, the alanine substitutions to more interior Cys/His residues (C38A, H63A, and C67A) result in strong repression that is no longer relieved by Ni.

**FIG 5.**
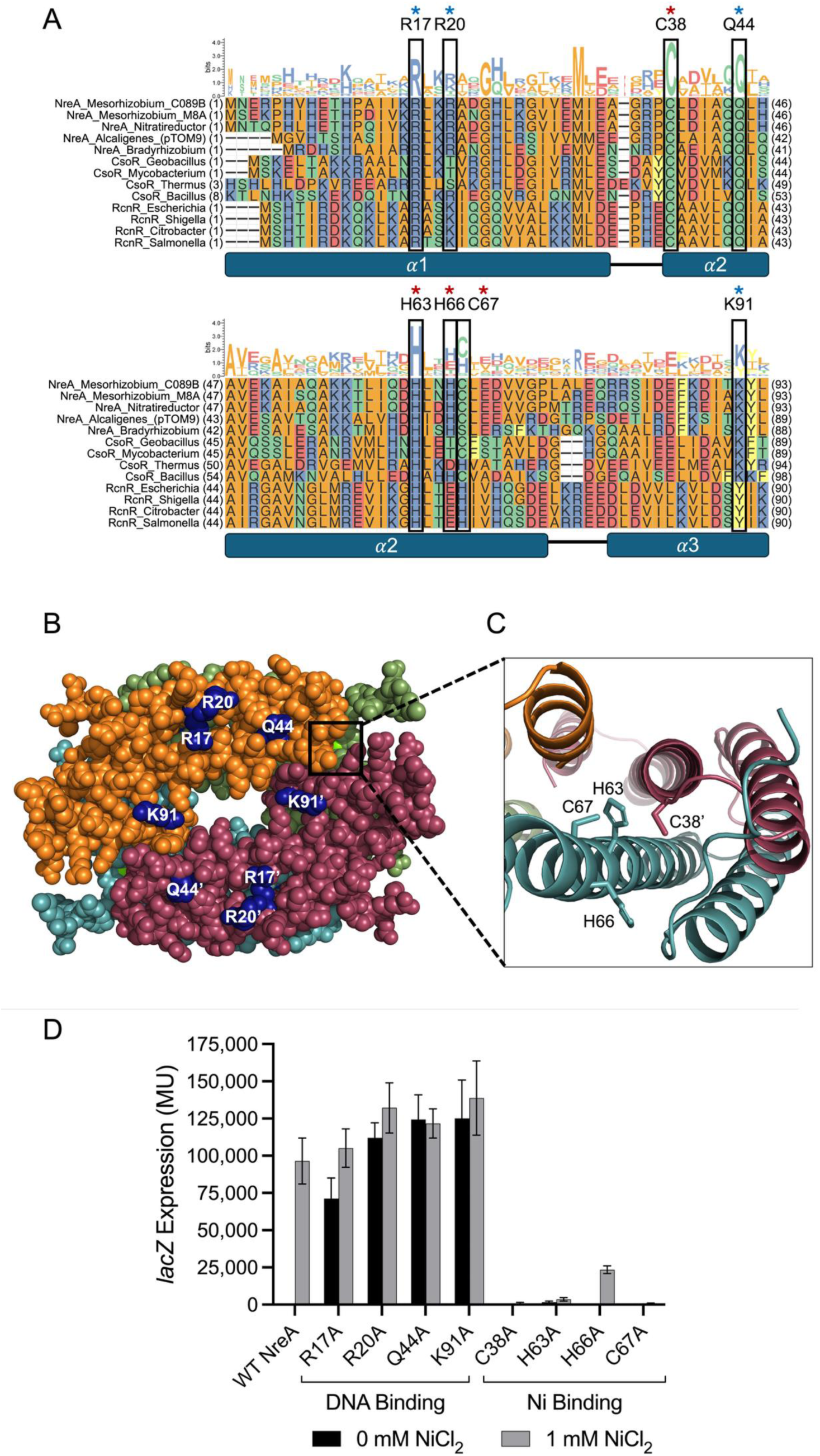
Influence of conserved NreA residues on NreA-mediated transcriptional regulation. (A) Protein alignment of NreA and CsoR/RcnR family members from diverse bacteria. Boxes indicate conserved residues that were individually altered in this experiment. Blue asterisks denote residues thought to be involved in DNA binding, while red asterisks denote residues thought to be involved in metal binding. (B) Alphafold3 prediction of the metal-bound form of NreA with proposed conserved DNA-interacting residues highlighted in blue. One of the four metal ions in this model is partially visible and indicated with a box. (C) A close-in view of the modeled metal-binding pocket. The metal ion has been removed from this rendering for clarity. (D) Reporter gene activity resulting from amino acid substitutions of NreA. *A. fabrum* cells harboring *lacZ* fusions to the −39GOF promoter/operator were grown in the presence or absence of supplemental NiCl_2_. β-galactosidase activity was measured in Miller units (MU). Error bars represent standard deviation from the mean (n=6).

## DISCUSSION

We have characterized the transcriptional control of a metal-responsive operon (*nreAXY*) in *Mesorhizobium* initially discovered for its role in evolutionary adaptation to Ni-rich serpentine soils (17, 19). Using phylogenetic analysis coupled with a heterologous *Agrobacterium*-based reporter system, we have identified NreA (a CsoR/RcnR family member) as a Ni-responsive transcriptional repressor, as well as *cis*-acting DNA sequences crucial to metalloregulation of this operon. We have further implicated specific residues in the NreA repressor as being important for either DNA binding or metal-mediated transcriptional induction.

Naturally diverse alleles of the *nre* operon in *Mesorhizobium* allowed us to pinpoint *cis*-acting regulatory sequences under strong selective pressure, and we mapped this *cis*-regulatory region to a remarkably compact segment of DNA comprising the 39 base pairs upstream of the *nreA* start codon. A partially palindromic operator sequence positioned within the −35/-10 spacer segment mediates repression by NreA. Operators interacting with CsoR/RcnR repressors are divided into two general types: Type 1 operators consist of a central GC-tract flanked by palindromic repeats, while Type 2 operators consist of a longer central region with two or more G/C-tracts flanked by palindromic repeats. In some CsoR/RcnR-controlled promoters, two operator sequences (of either type) may be situated in close proximity with each other (15, 18). The NreA operator (TATCCtctcgggggGGATA) is Type-1, with a single G_7_-tract. The palindromic repeats flanking the central G-rich region are critical for NreA-mediated repression (Fig. 4). Mutations to the G-rich central region result in more subtle de-repression phenotypes. However, operator variants designed to increase palindromic symmetry or to reduce G-richness while maintaining the consensus sequence, retained repression with some yielding stronger repression than the wild-type operator (Fig. S1). These findings suggest that NreA may function independently of DNA structural features such as A/B-form hybrids, which have been proposed to facilitate RcnR binding to G-rich operators (15, 18, 20). Indeed, NreA-interacting operator variants with a central GG→CA mutation showed enhanced repression, indicating that specific base pair contacts, rather than overall helical structure, may dominate NreA-DNA interactions (see Fig. S1).

Like other CsoR/RcnR family members, NreA monomers are small, at only 93 aa in length. While proteins in this family form alpha-helical bundles as monomers, they tend to form higher-order tetrameric structures (dimer-of-dimers) (9, 15, 18, 20–22). A structural model for NreA in a metal-bound state was generated by AlphaFold3 and this model places a metal-binding pocket with highly conserved His and Cys residues at intersubunit junctions (Fig. 5C) (23). Alanine substitution of the putative metal-coordinating residues (C38, H63, C67) resulted in a super-repressed state in our reporter assay system, consistent with failure to respond to Ni inducer (Fig. 5D). None of these metal binding pocket substitutions mimicked a metal-bound (promoter-on) phenotype; this may be attributed to the NreA tetramer being in a relaxed, DNA-binding state in the apo form, with metal binding introducing tension and DNA release (15). Substitutions containing side chains that are more bulky or positively charged may lead to metal-mimicking, non-repressing NreA variants. Only the tetrameric (or larger) configuration of NreA forms a modeled structure with the potential to bind metals and the palindromic operator of the length we have determined in this work. Docking of the NreA operator to the tetramer (using AlphaFold3) confirms a generally good fit of a positively charged tetramer surface to operator base pairs crucial for functionality (Fig. S2). Using structure-guided substitutions, we show that NreA-mediated repression is dependent on a set of conserved residues likely located on this DNA-binding surface (Fig. 5B), as alanine substitution of putative DNA-binding residues (R17, R20, Q44, or K91) led to constitutive promoter activity.

While considerable polymorphism exists among *nre*-containing *Mesorhizobium* strains–even in the *nreAXY* promoter region–they appear to uniformly produce a leaderless mRNA transcript devoid of a Shine-Dalgarno (SD) translation initiation enhancer, suggesting an adaptive advantage associated with this unusual mRNA structure. An apparent association between leaderless transcription and stress responses has previously been discussed (24, 25). When leaderless mRNAs engage directly with 70S ribosomes, certain time- and energy-consuming steps associated with conventional SD-mediated translation initiation are bypassed; in stressful circumstances, this enhanced economy may support higher levels of stress-response protein expression.

From an applied perspective, the NreA-controlled promoter/operator, with some engineering, is functional in *A. fabrum* and *E. coli*, demonstrating its broad portability and independence from species-specific factors. The full regulatory module (promoter/operator and regulator coding sequence) can be encoded on as little as 330 bp of DNA while enabling strongly inducible transgene expression. This genetic module is amenable to engineering for tuning the strength of repression or induction and possibly shifting metal specificity. Such a module could be adapted for use as a biosensor of various metals of biological and environmental importance. Super-repressor variants of NreA may be useful as compact DNA-binding reagents for applications such as in-vivo locus tracking or plasmid display.

## MATERIALS AND METHODS

### Bacterial strains and culture conditions

Strains used were *Mesorhizobium* C089B (17), *Agrobacterium fabrum* D224 (a streptomycin-resistant variant of (UBAPF2) (26), *Escherichia coli* DH5α (27) and DH5α-based conjugation helper strain B001 (28). Cells were cultured in Luria broth (LB; 10 g/L tryptone, 5 g/L yeast extract, 5 g/L NaCl_2_). When appropriate, 12 g/L of agar was added as a solidifying agent. Media supplements and antibiotics were as follows: streptomycin (Sm, 200 μg/mL), neomycin (Nm: 100 μg/mL), chloramphenicol (Cm: 30 μg/mL), and Nickel (II) chloride (1 mM). *Mesorhizobium* and *Agrobacterium* strains were grown at 30°C, and *E. coli* strains were grown at 37°C. Triparental matings were carried out at 30°C. All engineered strains used in this study are listed in Table S1 in the supplemental materials.

### Plasmid and strain construction

For testing promoter/operator constructs in *A. fabrum*, *lacZ* reporter plasmids pKJ138 (*nreA*-) and pKJ140 (*nreA*+) were used (Fig. 2B, sequences in Supplemental Appendix 1). Promoter/operator variants (assembled as hybridized oligonucleotides) were ligated upstream of *lacZ* using BamHI and XbaI sites (Table 1). For creating NreA alanine substitution variants (Table 2), the wild-type *nreA* gene was first cloned in the small plasmid pKJ213 (Fig. S3, sequence in Supplemental Appendix 1), with which mutagenesis oligonucleotides were used to incorporate changes via round-the-horn PCR followed by circularization with T4 DNA ligase and T4 polynucleotide kinase. The resulting *nreA* variants were then amplified with primers oKJ614 and oKJ615 and inserted into pKJ227 (pKJ138 with −39GOF promoter/operator), via a KpnI site. All oligonucleotide sequences used for producing promoter/operator and *nreA* variants are given in Table S2 in the supplemental materials. All plasmids used in this study were verified by Sanger sequencing. Reporter plasmids were transferred to *A. fabrum* D224 via triparental mating using the helper strain (B001) and donor strains (DH5α) containing the experimental plasmids. For conjugation, strain mixtures were co-cultured on LB agar at 30°C for 4 to 24 h and plated on LB agar containing Sm and Nm.

**Table 1.**
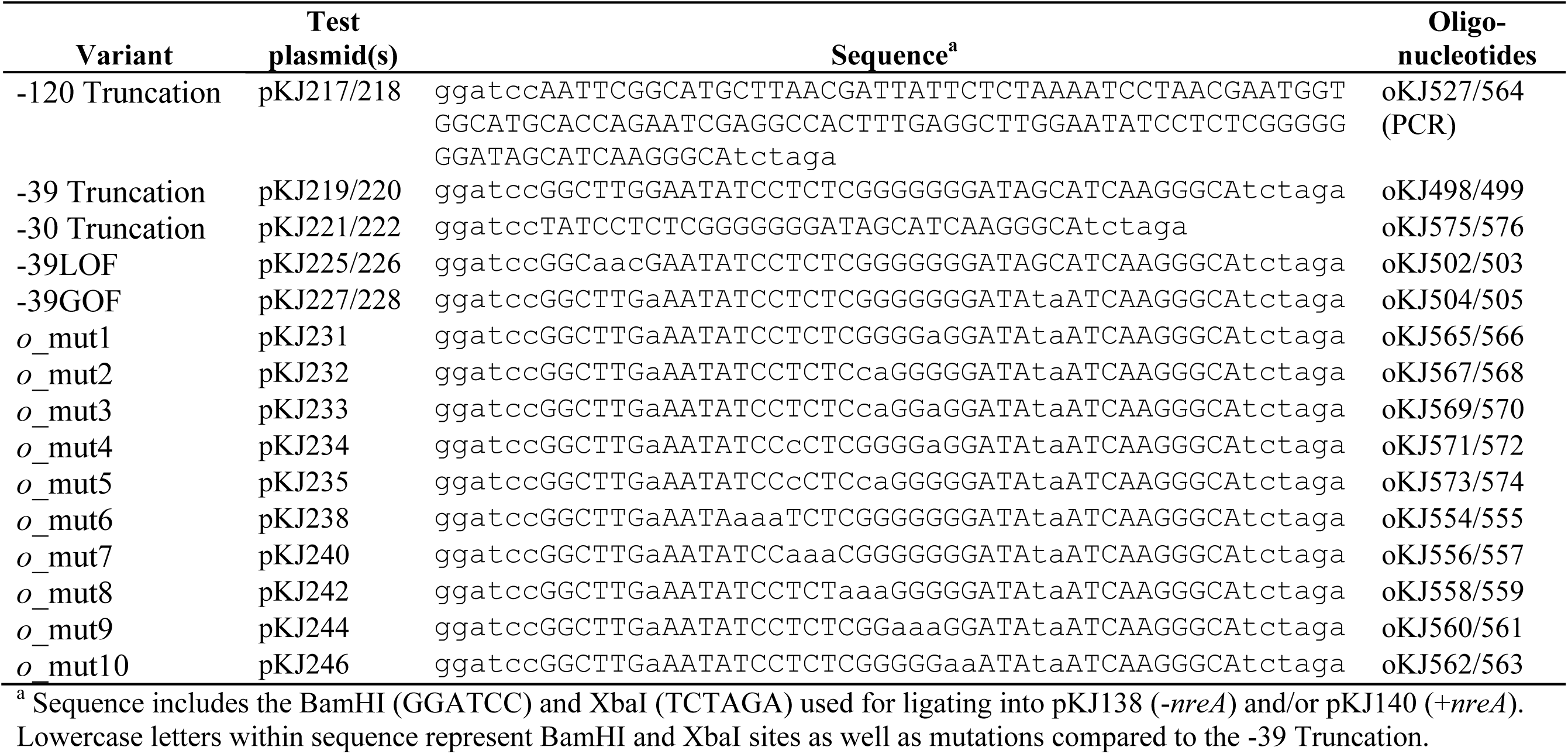
Variant promoter/operator sequences in this study.

**Table 2.**
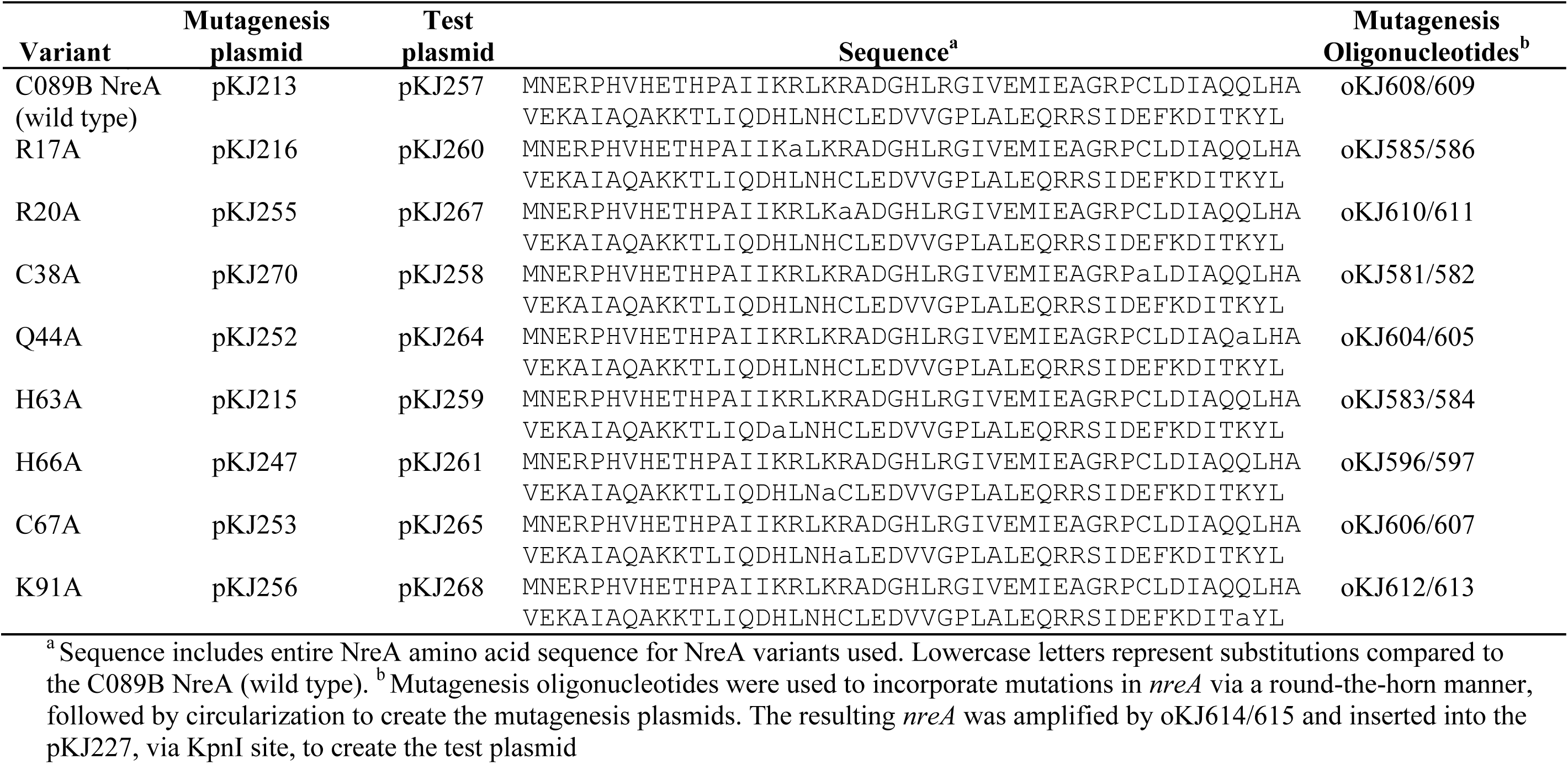
NreA variants used in this study.

#### *lacZ* expression assays

Modified Miller assays similar to [Zhang and Bremer (29)] were carried out as follows: Bacteria were grown in liquid culture overnight and their absorbance at 600 nm (A_600_) was measured; culture density was then adjusted to an A_600_ value of 0.2 using LB broth. These suspensions were diluted 10-fold into 200 uL LB in 96-well plates and cultured with shaking (∼300 rpm) at 30°C for 6 h, with half-strength antibiotics (Sm-100 and Nm-50). From these cultures, 2 uL of suspension was transferred to 23 µL of permeabilization solution (100 mM Na_2_HPO_4_, 20 mM KCl, 2 mM MgSO_4_, 0.8 mg/mL hexadecyltrimethylammonium bromide, 0.4 mg/mL sodium deoxycholate, 5.4 µL/mL beta-mercaptoethanol), incubated at 30°C on a shaker for 5 min, after which 150 µL of Substrate solution (60 mM Na_2_HPO_4_, 40 mM NaH_2_PO_4_, 1 mg/mL o-nitrophenyl-β-D-galactoside, 2.7 µL/mL beta-mercaptoethanol) was added to each well. After sufficient yellowing occurred (around 30 min) 175 µL of stop solution (1 M Na_2_CO_3_) was added to each well and their absorbance at 420 nm (A_420_) was measured. For detection of very low *lacZ* expression (Fig. S1), overnight cultures were normalized to an A_600_ value of 2.0, and 5 µL of these suspensions were used in Miller assays as described above, with a reaction time of approximately 90 min prior to adding stop buffer.

### Transcription start site mapping (5’-RACE)

The *A. fabrum* strain, harboring pKJ218, was cultured in LB supplemented with half-strength Sm Nm and 1 mM NiCl_2_ to an A_600_ of ∼0.8. A cell pellet was then made from 1.5 mL of culture and this was placed at −80°C for 15 min. The frozen pellet was thawed and resuspended in 50 µL of cold TED solution (20 mM Tris (pH 7.5); 10 mM EDTA (pH 8.0); 40 mM dithiothreitol) and 50 µL of 1% SDS. The resuspended pellet was vortexed briefly and immediately placed on a 70°C heat block for 3 min. 1 mL of Tri-reagent (Trizol) was added to the suspension, inverting vigorously. After ensuring the solution was thoroughly mixed, 200 µL of chloroform was added, vortexed briefly, and the suspension was centrifuged for 2 min at 15,000 rpm. After centrifugation the aqueous layer was separated and placed on ice. 600 µL of isopropanol was then added to the aqueous layer to precipitate the RNA. The RNA suspension was kept on ice for 5 min and then centrifuged. After centrifugation, supernatant was removed and the pellet was then resuspended in 40 µL of RNAse-free water and placed in a 70°C heat block. Approximately 1 µg of RNA was reverse transcribed into cDNA using 6 µM gene-specific primer oKJ591, 5x ProtoScript II Buffer, 0.5 mM dNTPs, 10 µM DTT and 200 U of ProtoScript II Reverse Transcriptase (NEB M0368). The mixture was incubated at 42°C for 1 hr. After incubation, the product was purified using a Zymo ZR^TM^ Plasmid Miniprep-Classic kit (D4016), and the cDNA was eluted in 30 µL of T_3_E_0.3_. Polyadenylate [poly(A)] was added to the 3’ ends of the cDNA molecules using 10x terminal transferase (TdT) buffer, 0.1 mM dATP, 0.25 mM CoCl_2_, and 15 U of TdT (NEB M0315). This was incubated at 37°C for 30 min. The product was placed into a 70°C water bath for 10 min for heat inactivation. The heat inactivated product was amplified using Taq polymerase (NEB M0267), a gene-specific primer oKJ591, and oKJ595 (this primer consists of a (dT)_18_ at its 3’-end and a G/C rich adapter sequence at its 5’-end). The amplification protocol included 94°C for 2 min, followed by 10 cycles of 94°C for 15 s, 39°C for 20 s, and 70°C for 45 s, followed by 28 cycles of 94°C for 15 s, 55°C for 20 s, and 70°C for 45 s, and a final extension step at 70°C for 2 min. The product was used in a second round of PCR using a nested primer, oKJ592, along with the (dT)_18_ primer oKJ595. The amplification protocol included 94°C for 2 min, followed by 36 cycles of 94°C for 15 s, 55°C for 20 s, and 70°C for 45 s, and a final extension step at 70°C for 2 min. The product was used in a third round of PCR using a nested primer, oKJ593, along with the (dT)_18_ primer oKJ595. The amplification protocol included 94°C for 2 min, followed by 34 cycles of 94°C for 15 s, 56°C for 20 s, and 70°C for 45 s, and a final extension step at 70°C for 2 min. The product was then cleaned up using a Zymo ZR^TM^ Plasmid Miniprep-Classic kit (D4016) and submitted for Sanger sequencing using the primer oKJ593.

### Sequence alignments and phylogenetic analysis

An alignment was performed with nucleotide sequences of the region upstream of the *nreAXY* operon acquired from various *Mesorhizobium* strains. Strains and sequences used are listed in Table S3. The sequences were aligned using (clustalW) and the figure was created in R using ggmsa and WebLogo3 (30, 31). An alignment of NreA/CsoR/RcnR protein sequences (listed in Table S4) was performed using clustalW and the figure was created on R using ggmsa (30). A phylogenetic tree reconstruction was built using the Phylogeny.fr analytical server. This online tool used an existing ClustalW-generated multiple sequence alignment of 13 representative amino acid sequences (above) and re-aligned them using MUSCLE. It then used Gblocks with the highest-stringency settings, which minimizes contiguous strings of non-conserved residues.

PhyML (v3.1/3.0 aLRT) was then used to reconstruct a maximum likelihood-based phylogenetic tree with 100 bootstrap replicates. The WAG substitution model was selected assuming an estimated proportion of invariant sites (of 0.116) and four gamma-distributed rate categories to account for rate heterogeneity across sites. The gamma shape parameter was estimated directly from the data (gamma=3.991). Reliability for internal branch was assessed using the aLRT test (SH-Like). TreeDyn was used to visualize the tree reconstruction (32–38).

## ACKNOWLEDGMENTS

This study was supported by the National Science Foundation (NSF) and by the Department of Microbiology and Molecular Biology at Brigham Young University

## AUTHOR CONTRIBUTIONS

Kyson T. Jensen^a^, experimental design, experimental analysis, writing | Joel S. Griffitts^a^, funding acquisition, project administration, experimental design, writing

## DATA AVAILABILITY

All data is available upon request.

## ADDITIONAL FILES

### Supplemental Material

FIG S1—FIG S3; Table S1—Table S4; Sequences for plasmids pKJ138, pKJ140, and pKJ213.

